# Variation in phenotypic plasticity in desiccation tolerance is driven by trade-offs, not climate, in *Drosophila melanogaster*

**DOI:** 10.1101/2025.05.12.653617

**Authors:** V Kellermann, B van Heerwaarden

## Abstract

The capacity for species to respond to environmental variation via phenotypic plasticity has been proposed as a mechanism for buffering species against climate change. Two main theories are proposed to explain the evolution of phenotypic plasticity – the climate variability hypothesis and the trade-off hypothesis - but the evidence for these hypotheses remains mixed. In the current study, we examine phenotypic plasticity (hardening capacity) in desiccation resistance across populations of *Drosophila melanogaster* collected along a climatic gradient from eastern Australia. While climate was an important driver of innate desiccation tolerance, we found no evidence of climate shaping hardened desiccation tolerance or hardening capacity. Instead, we found that populations with high tolerance tended to have lower plasticity, indicative of a trade-off. Further analyses, accounting for statistical non-independence, showed a strong negative correlation between innate desiccation tolerance and hardening capacity, supporting the trade-off hypothesis. These results suggest that for species with high tolerance, phenotypic plasticity is unlikely to contribute to their response to climate change.

## INTRODUCTION

How species respond to climate change will depend, in part, on their capacity for evolutionary shifts in key traits. Over longer time frames, this is likely to involve genetic change, i.e. changes in allele frequencies, but in the short term, species may respond/adapt via phenotypic plasticity, here defined as a rapid within-generation response to the environment (Chevin et al. 2010, Chevin and Hoffmann 2017). Plastic increases in tolerance can be induced through short-term (minutes/hours) hardening or longer-term (days/weeks) acclimation treatments. In ectotherms, climatic resilience traits like heat and desiccation tolerance (surviving low humidity) have been linked to species distributions and range limits (Chown et al. 2011, Kellermann and van Heerwaarden 2019). Consequently, the capacity for species to shift these traits via evolutionary processes (genetic adaptation and plasticity) is likely to underpin climate change responses (Chevin et al. 2010, Schilthuizen and Kellermann 2014, Sgro et al. 2016). However, the extent to which phenotypic plasticity is under selection and, therefore, evolves in response to different environments is still largely unknown (Ghalambor et al. 2006, Sgro et al. 2016).

Two key hypotheses have been proposed to explain why species and populations may vary in their capacity to respond to environmental variation via phenotypic plasticity: the tolerance-plasticity trade-off hypothesis and the climatic variability hypothesis. The tolerance-plasticity trade-off hypothesis predicts that a trade-off between innate tolerance and plasticity constrains the evolution of high tolerance and high plasticity (Stillman 2003, Gunderson and Stillman 2015, van Heerwaarden and Kellermann 2020, Barley et al. 2021). That is, individuals can have high tolerance or high phenotypic plasticity but not both. Trade-offs between tolerance and plasticity across populations or species are often used to imply a genetic or physiological constraint in the evolution of tolerance traits. Genetic constraints arise when the same genes that underpin tolerance and plasticity have opposing effects on fitness (Roff and Fairbairn 2007, van Heerwaarden and Kellermann 2020). Resource acquisition–allocation theory, primarily developed in the context of life-history evolution, suggests that organisms face resource limitations and must allocate those resources strategically to maximise fitness ()(van Noordwijk and de Jong 1986, Roff and Fairbairn 2007). Under this framework, investment in one trait, such as thermal tolerance, may come at the cost of reduced plasticity. Physiological limits can also drive trade-offs between tolerance and plasticity because species that are close to their physiological limit will have limited capacity to shift traits via plasticity (Sorensen et al. 2016). Notably, there are also methodological reasons, unrelated to constraints, why trade-offs may emerge in datasets that should also be considered (van Heerwaarden and Kellermann 2020).

Empirical evidence for the tolerance-plasticity trade-off hypothesis is mixed (Gunderson and Stillman 2015, van Heerwaarden et al. 2016). However, the tolerance-plasticity trade-off hypothesis has been plagued by limitations linked to how the trade-off is analysed (van Heerwaarden and Kellermann 2020, Gunderson 2023), and how phenotypic plasticity is measured/calculated. Many studies use only a small number of acclimation or hardening treatments (Barley et al. 2021), which may underestimate plastic responses or entirely miss the treatments that may induce a plastic response, particularly those with high innate tolerance (van Heerwaarden et al. 2024). In *Drosophila*, species with higher tolerance required longer hardening treatments to induce plastic responses (threshold shift hypothesis) for both desiccation and heat tolerance (van Heerwaarden et al. 2024). Trade-offs may emerge in datasets if more extreme treatments that induce a plastic response in species with higher tolerance are not included in the design. Statistical artifacts also manifest, particularly when only a handful of plasticity treatments are used (Kelly and Price 2005, Gunderson 2023). This is because plasticity is typically calculated by taking the difference between temperatures/treatments and innate tolerance (which is taken from one of these environments), meaning the comparison between innate tolerance and plasticity defies statistical independence (van Heerwaarden and Kellermann 2020). These issues can be resolved by using multiple hardening treatments or correcting for the statistical artefact when estimating plasticity. However, current trade-off datasets rarely account for these issues, meaning it is difficult to confirm whether there is a trade-off between plasticity and tolerance (Gunderson 2023).

An alternate, but not necessarily mutually exclusive, hypothesis is the climatic variability hypothesis. This hypothesis predicts species/populations occupying more variable environments should evolve higher phenotypic plasticity to cope with environmental fluctuations (Janzen 1967, Ghalambor et al. 2006). A key component of this hypothesis is that environmental variability is somewhat predictable to counter the costs of a mismatch in phenotypes if environments change unpredictably because the costs to plasticity are generally high (Gabriel and Lynch 1992). The results of inter and intra-specific studies in ectotherms, predominately thermal tolerance traits, have found equivocal support for the climatic variability hypothesis (Gunderson and Stillman 2015, Barley et al. 2021). Many of these studies, however, examined only a handful of populations, making it difficult to establish whether an absence of support for the climatic variability hypothesis was because studies lacked the power to detect associations between plastic capacity and environment. Furthermore, the climatic variability hypothesis can suffer from similar issues to the trade-off hypothesis; that is, if there are threshold shifts and only a handful of treatments are used and the full range of plastic responses are not captured, we may underestimate plastic responses in species/populations with high innate tolerance, those that typically occupy more variable environments.

A more powerful approach to study the climatic variability hypothesis is to examine multiple populations along a latitudinal cline/climatic gradient. Clinal studies have revealed genetic divergence and plasticity in numerous life history and tolerance traits across populations (Hoffmann et al. 2003, Sinclair et al. 2012) and have been instrumental in identifying key traits that are likely under climatic selection. Some studies have tested the climatic variability hypothesis along a cline. Studies on copepods and the Mediterranean fruit fly found no evidence supporting the climatic variability hypothesis in tolerance traits (CTmin/CTmax/desiccation and starvation tolerance) (Pereira et al. 2017, Weldon et al. 2018). Similar patterns were also absent in *Drosophila simulans*, where plasticity in both cold and heat tolerance was not associated with latitude/climate (van Heerwaarden et al. 2014). Clinal variation in CTmax and CTmin were found in a collembolan, but only clinal patterns in plasticity for CTmax were found (Jensen et al. 2019). Significant clinal patterns were found in heat hardening in *D. melanogaster* but in the opposite direction expected based on the climatic variability hypothesis (Sgro et al. 2010), such that tropical populations were more plastic than the temperate populations, although this pattern was weak. Clinal studies of phenotypic plasticity have consistently found population-by-treatment interactions, indicating genetic divergence between populations and supporting the idea that genetic variation underpins phenotypic plasticity. But inconsistent relationships between plastic capacity and climate/latitude suggest plastic capacity is not always under climatic selection. It is also possible that methodological limitations outlined above may also contribute to these mixed patterns.

In the current study, we simultaneously explored both plasticity hypotheses (trade-off/climatic variability hypothesis) in desiccation tolerance, a trait important in shaping the distribution of terrestrial arthropods, including *Drosophila* (Kellermann et al. 2012). Phenotypic plasticity for desiccation tolerance, measured as rapid hardening, has also been shown in several *Drosophila* species, including *D. melanogaster* (Kellermann et al. 2018). In *Drosophila melanogaster*, innate desiccation tolerance shows latitudinal clinal variation in Australia, with populations from temperate populations more tolerant to desiccation stress than tropical populations (Lasne et al. 2018). A study on a low and high-latitude population of *D. melanogaster* found lower plasticity in desiccation tolerance in the high-latitude (temperate) population (Clemson et al. 2018) in opposition with the climatic variability hypothesis. However, more extensive sampling is needed to rule out whether climatic variability drives the evolution of desiccation plasticity in this species. The high latitude population also had the highest innate desiccation tolerance, in line with the trade-off hypothesis. However, again this was based on only two populations and used one hardening treatment, which may not have been strong enough for the temperate populations (if there are threshold shifts).

Leveraging the climatic gradient along the east-coast Australia, we examined clinal variation in innate desiccation tolerance and hardening capacity in ten populations of *D. melanogaster*. In a smaller subset of populations, we first asked to what extent different hardening treatments influence plastic responses and whether thresholds underlie phenotypic plasticity. Across the entire ten populations, we then asked whether there is evidence for genetic differentiation across populations in their plastic response and whether population variation in phenotypic plasticity is associated with different climate measures. Finally, we explored whether there is evidence for a trade-off between innate desiccation tolerance and hardening capacity.

## MATERIALS AND METHODS

Populations of *Drosophila melanogaster* were collected from 10 locations along the east coast of Australia between March and April 2012 (Fig. 1, Table S1). Approximately 20 field-inseminated females were collected from each site and used to establish isofemale lines for each population. Following three generations of laboratory culture, approximately 10 males and 10 females from each isofemale line were used to generate a mass-bred population for each collection site.

**Figure 1.**
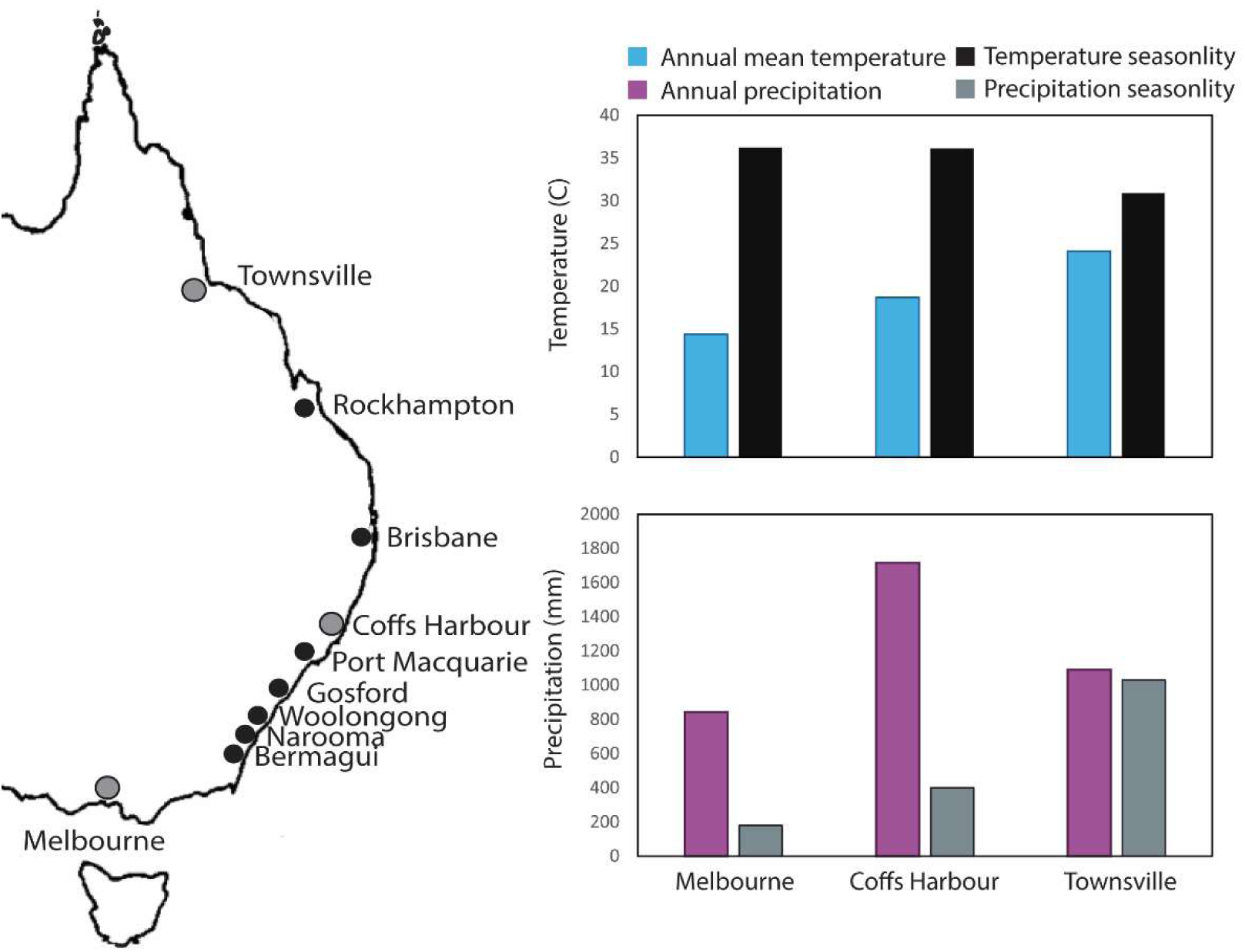
Populations of *D. melanogaster* collected along a latitudinal and climatic gradient along the east-coast of Australia and temperature variables: annual mean temperature and temperature seasonality, as well as precipitation variables: annual precipitation and precipitation seasonality for three populations collected along the latitudinal cline.

Each mass-bred population was kept at 25°C under a 12:12 light/dark cycle in two 250ml bottles containing 60mL of a potato, yeast and dextrose media (37.32 % yeast, 31.91 % dextrose, 23.40 % potato medium and 7.45 % agar combined with 98.48 % H2O, 0.97 % ethanol, 0.45 % propionic acid and 0.11 % nipagen). Flies were transferred to fresh bottles every two to three days. Populations were kept at a density of 300-400 flies per bottle to ensure census populations of approximately 800 flies for each collection site. These populations were maintained for nine generations in the laboratory prior to experiments.

One generation before the experiment was initiated, each mass-bred population was transferred into 8x8cm laying pots. A thin layer of live yeast paste was also spread over the media to encourage oviposition. Each laying pot contained approximately 100-200 flies and was left for 20 hours at 25°C. The eggs from these laying pots were collected and transferred to 40mL vials containing 6mL of media. A total of nine vials per population were collected, with each vial containing a density of approximately 50 eggs, as larval density has been shown to influence estimates of fitness and stress traits (Sorensen and Loeschcke 2001). The focal offspring were collected from these vials within one day of eclosing and held in fresh vials containing 4mL of media. The flies were left for three to four days to ensure all females were mated. Following this, females and males were separated under CO2 anaesthesia. Flies were given a 2-day recovery period to avoid any adverse consequences of CO_2_ exposure on the estimation of traits.

### Threshold shifts amongst populations

To assess the capacity of different populations to respond to a desiccation hardening treatment, and whether populations differed in the threshold needed to induce a hardening response, we first exposed flies from three populations to a series of desiccation pre-treatments. This pre-treatment involved placing flies in groups of 10 into empty vials (dimensions) with no food or water source (food contains moisture and therefore cannot be added to the hardening regime) and placing them into a tank with 5-10% RH. Because previous experiments have shown that species may vary in the exposure times needed to induce a hardening response and this could depend on innate tolerance (i.e. they vary in their hardening threshold) (van Heerwaarden et al 2024), we examined four exposure times that were most likely to induce a hardening response without causing mortality. If the desiccation pre-treatment causes mortality, this would essentially be imposing selection, which would artificially shift the mean higher as the least resistant individuals will have died. We chose four exposure periods based on a-priori knowledge of desiccation tolerance in this species, 9, 10, 11 and 12hrs. Following these exposure periods, we gave flies a 9hr recovery period on a food and water source. This recovery period was chosen based on pilot experiments, where exposures over 12hrs tended to induce mortality. The exposure periods were initiated such that all flies were moved from the pre-treatment to the recovery treatment at the same time. At the same time as setting up the hardening pre-treatment control flies were also placed into groups of 10 but were given a food source containing moisture. On completion of the recovery period, individual flies were moved into vials which were then placed into a tank at 5-10 RH%. Flies were scored every hour until all flies had succumbed to desiccation.

### Clinal study

We then tested for evidence of clinal variation in innate and hardening desiccation tolerance. We chose the desiccation hardening treatment time based on the threshold experiments and using the same methods described above, we exposed all ten populations to an 11-hour desiccation stress, followed by a 9-hour recovery period, at which point flies were placed back into the tank and scored for desiccation stress. For each treatment and population, 20 flies were assessed.

## STATISTICAL ANALYSIS

### Threshold shift

To test whether populations varied in how they responded to the hardening treatment, we analysed the threshold shift experiment with a general-linear model with Type III sums of squares in R. We included desiccation hardening treatment and population as fixed effects and tested for the interaction between the two (population variation in hardening response). Using this data we also calculated the hardening response ratio (modified from Gunderson and Stillman 2015), to look at trade-offs between traits. The hardening response ratio was calculated as (innate tolerance – hardened tolerance)/Treatment Time(hrs). We calculated this for each treatment for each population.

#### Environmental data

Environmental data was extracted from the WorldClim dataset based on the GPS coordinates from the collection location in R (Team 2014). Because there is significant auto-correlation between many of the environmental variables, we focused on ones we know to be linked to desiccation tolerance and desiccation plasticity in the past (Kellermann et al. 2012) – maximum temperature of the environment (T_MAX_), minimum temperature of the environment (T_MIN_) and annual precipitation (P_ANN_) (note: annual mean temperature is highly auto-correlated with T_MIN_), and because we were examining plasticity-related traits, we also included temperature seasonality and precipitation seasonality variables. Latitude is traditionally used in clinal studies as a proxy for climate. Consequently, we included latitude as an alternative predictor variable.

#### Plasticity metrics

When estimating the plastic response in any trait, you capture both the plastic and innate responses. To examine how plasticity differs across populations, it can be useful to look explicitly at the plastic component. To do this, we used a commonly used metric: hardening capacity (HC), which is calculated as hardened tolerance – innate tolerance. Hardening capacity shows the absolute difference between innate and hardened flies. Because hardening capacity and innate tolerance are often calculated from the same individuals and, therefore, not statistically independent from each other, we also divided our estimates of innate tolerance into two groups (reducing the replication of innate tolerance), one group was used to estimate innate tolerance and the other used to estimate hardening capacity, eliminating the issue of statistical independence.

#### Clinal study

We first asked whether populations varied in their innate and hardened desiccation tolerance by analysing the data with a general-linear model and analysis of variance (Type III). Population, treatment (innate vs hardened) and tank were treated as fixed factors in the model. We then asked whether innate and hardened tolerance varied in a way that suggests climatic selection acting on these traits. Here, we used a mixed-effects general linear model and analysis of variance in the R statistical package (Team 2014). Population was deemed a random effect, while treatment, tank and environmental variables including latitude (see above) were treated as fixed factors. We used the raw trait values for innate and hardened desiccation tolerance and our plasticity (hardening capacity) measure in all the models. Because we found a significant treatment x environment interaction, we separated the analysis by treatment to look at the strength of the relationship between basal desiccation tolerance and hardened desiccation tolerance and the environment independently.

## RESULTS

### Is there evidence for threshold shifts?

One reason we might see population differences in hardening responses is that populations with different innate tolerances require different exposure times to desiccation stress to elicit a response (threshold shift hypothesis, van Heerwaarden and Kellermann 2020). To test this, we examined whether desiccation hardening responses varied across three populations (northern, central and southern) of *D. melanogaster* exposed to varying durations of a desiccation stress pre-treatment (hardening). We found a significant difference between innate and hardened flies, i.e. hardening response (Table 1., Fig. 2) such that flies exposed to a pre-treatment desiccation stress were more desiccation resistant than flies not exposed to the pre-treatment. We also found a significant treatment x population interaction; this was most likely driven by the Coffs Harbour population, where hardening resulted in the largest increase in desiccation tolerance and the Townsville population which did not show a hardening response (Table 1., Fig. 2). When we looked solely at hardened flies, we found different exposure times did not significantly influence desiccation tolerance, with all hardening times having comparable levels of desiccation tolerance (Table 1., Fig. 2).

**Figure 2.**
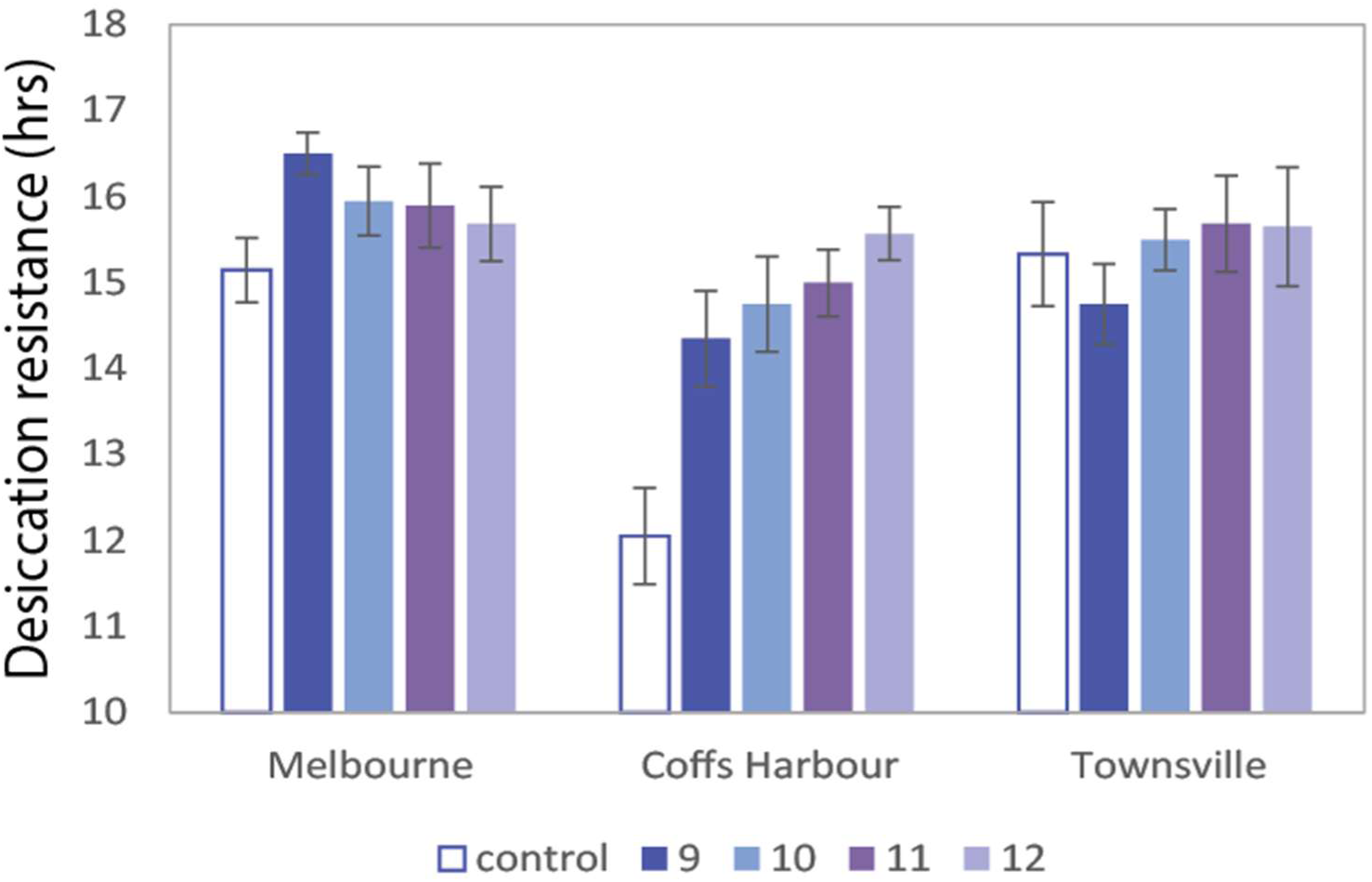
Average desiccation tolerance for basal and flies hardened for varying lengths of time for northern, central and southern populations of *D. melanogaster*.

**Table 1.** **A)** Analysis of variance examining whether desiccation tolerance varied between no exposure to pre-treatment of desiccation stress vs pre-treatment (Treatment) and whether this differed between three populations of *D. melanogaster.* **B)** Analysis of variance examining whether desiccation tolerance varied between the different exposure times (hardening duration) and whether the response to different exposure times varied across populations. Note that flies not treated with pre-treatment desiccation stress were not included in B).

| A) | D.F | F-value |
| --- | --- | --- |
| Treatment | 1 | 27.019*** |
| Population | 2 | 13.796*** |
| Treatment x Population | 2 | 6.884** |
| Residuals | 287 |  |

| B) | D.F | F-value |
| --- | --- | --- |
| Hardening duration | 3 | 1.227 |
| Population | 2 | 5.975** |
| Hardening duration x Population | 6 | 1.067 |
| Residuals | 225 |  |

### Do hardening responses vary across populations and climate?

Across all ten populations we found that an 11hr hardening treatment increased desiccation tolerance by an average of 2hrs across all populations. The capacity to harden varied across populations as evidenced by a significant population-by-treatment interaction (Population x Treatment: df=9, F-value = 2.956, P = 0.002), with hardening responses as little as a 40-minute increase in some populations and as great as 4 hours in others (Fig 3).

**Figure 3.**
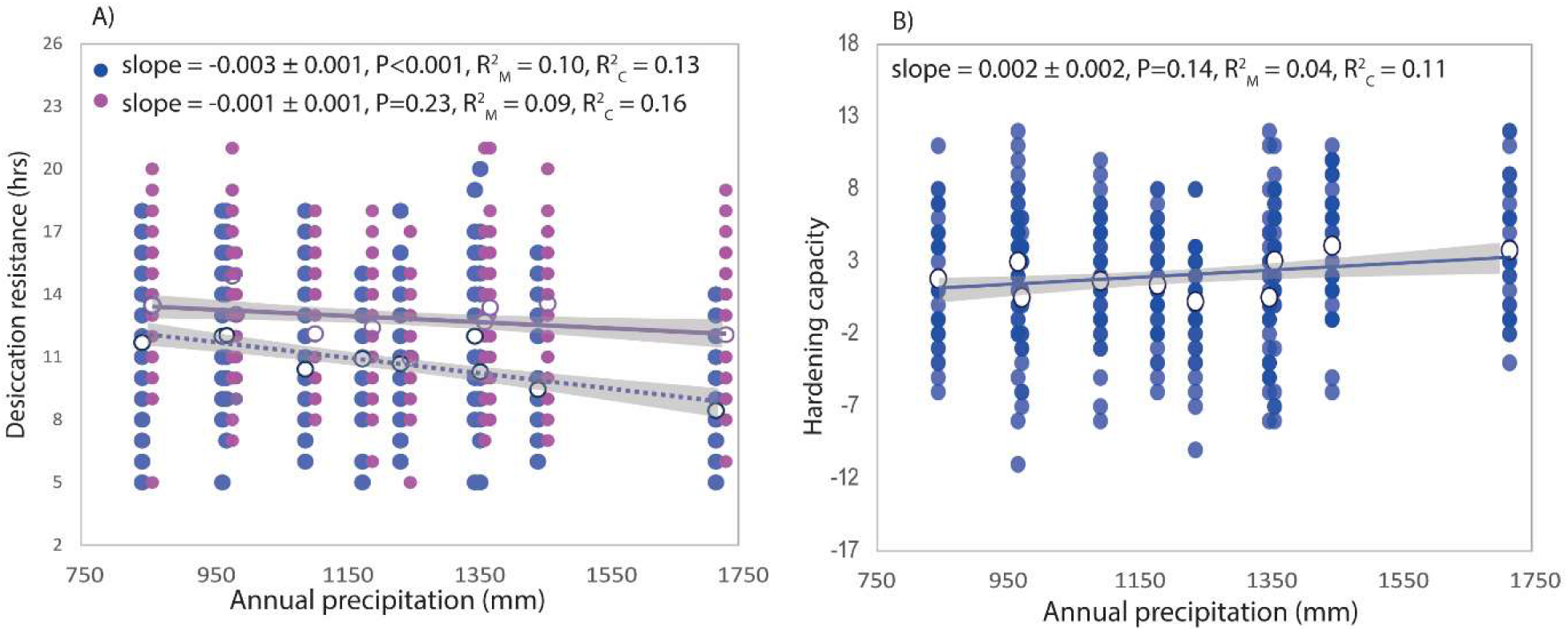
**A)** The relationship between desiccation tolerance and annual precipitation for hardened (purple dots) and basal (blue dots) desiccation tolerance in *D. melanogaster* populations. **B)** The relationship between hardening capacity and annual precipitation in *D. melanogaster* populations. The results from the linear model are presented, including the slope, and marginal R^2^_M_ (fixed effects) and conditional R^2^ (fixed and random effects).

Next, we asked whether populations varied in their innate and hardened desiccation tolerance, consistent with climatic selection acting on these traits, using all 10 populations. We looked at several climatic variables, including those related to variability (temperature and precipitation seasonality), and found that annual precipitation could explain the largest proportion of variation in our data, such that desiccation resistance increased with decreasing annual precipitation (Table 1). We did not find any climate seasonality measures or latitude that could explain variation in desiccation or hardened desiccation tolerance (Table S2 and 3, Fig S1). A significant interaction between treatment and annual precipitation was detected such that the relationship between annual precipitation was strongest for innate desiccation tolerance (Table 1, Fig 3A). Further dissecting the data to look at the strength of the relationship between annual precipitation and innate and hardened desiccation tolerance, we found a weak (R ^2^=0.10) but significant association for innate tolerance (Fig 3A), but no relationship was detected for hardened desiccation tolerance, suggesting that innate, but not hardened desiccation tolerance is under climatic selection (Fig 3A). When we examined the capacity for species to harden using hardening capacity, we found no relationship between hardening capacity and climate (Fig 3B).

**Table 1.**
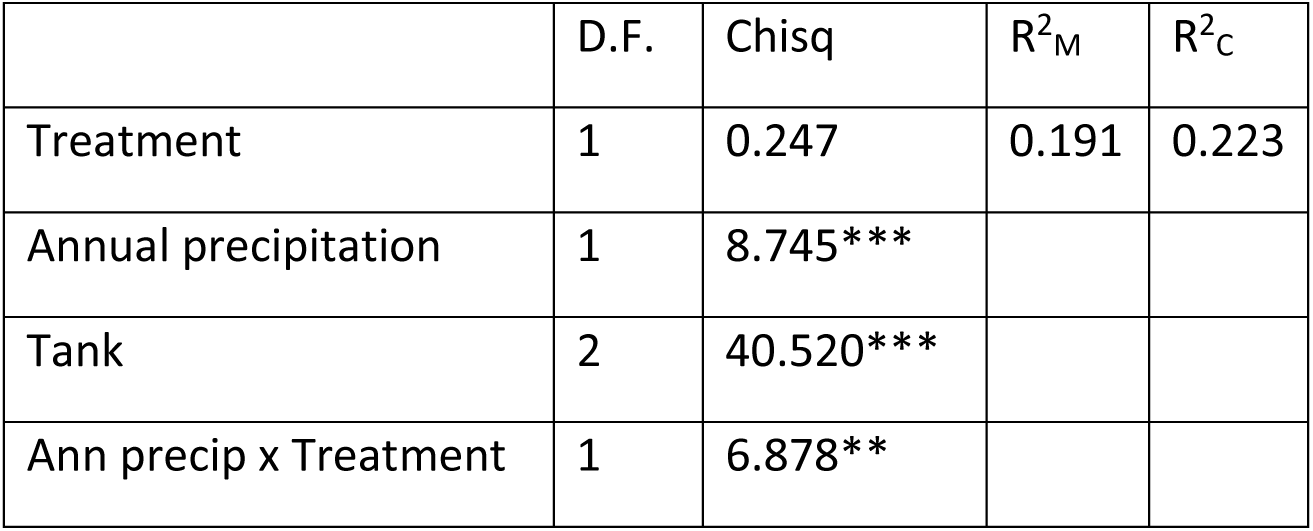
Mixed effects analysis of variance testing for an effect of treatment (control vs hardened) and precipitation (of collection site) on desiccation tolerance. The proportion of variation explained by the fixed effects (R^2^_M_) and random effects (R^2^_C_) are shown.

| | D.F. | Chisq | $R^2_M$ | $R^2_C$ |
| --- | --- | --- | --- | --- |
| Treatment | 1 | 0.247 | 0.191 | 0.223 |
| Annual precipitation | 1 | 8.745*** |  |  |
| Tank | 2 | 40.520*** |  |  |
| Ann precip x Treatment | 1 | 6.878** |  |  |

### Is there a trade-off between innate desiccation tolerance?

Finally, we looked for a trade-off between the capacity to respond via plasticity (hardening capacity) and innate tolerance. We first used a common method of looking for trade-offs by regressing hardening capacity against innate desiccation tolerance. Using this method, we found evidence for a substantial trade-off (Fig 4A). However, this analysis method can lack statistical independence (Kelly and Price 2005, van Heerwaarden and Kellermann 2020). To overcome this, we took two approaches: firstly, we created an independent estimate of innate tolerance and hardening capacity by splitting our data in half. Using half of the estimates, we averaged innate tolerance across all the populations, and with the remaining half, we estimated hardening capacity. With this method, the negative relationship between innate tolerance and hardening capacity remained (Fig. 4B). Finally, we went back to our data that captured hardening responses across multiple hardening treatments (for three populations) and calculated the hardening response ratio, thus reducing some of the statistical bias between innate tolerance and hardening capacity (Fig 4B). While we did not have the power to detect significant associations between innate tolerance and plastic capacity with only three populations, relating hardening response ratio to innate tolerance showed a clear trend that populations with higher tolerance tended to have a smaller plastic response (Fig 4B).

**Figure 4.**
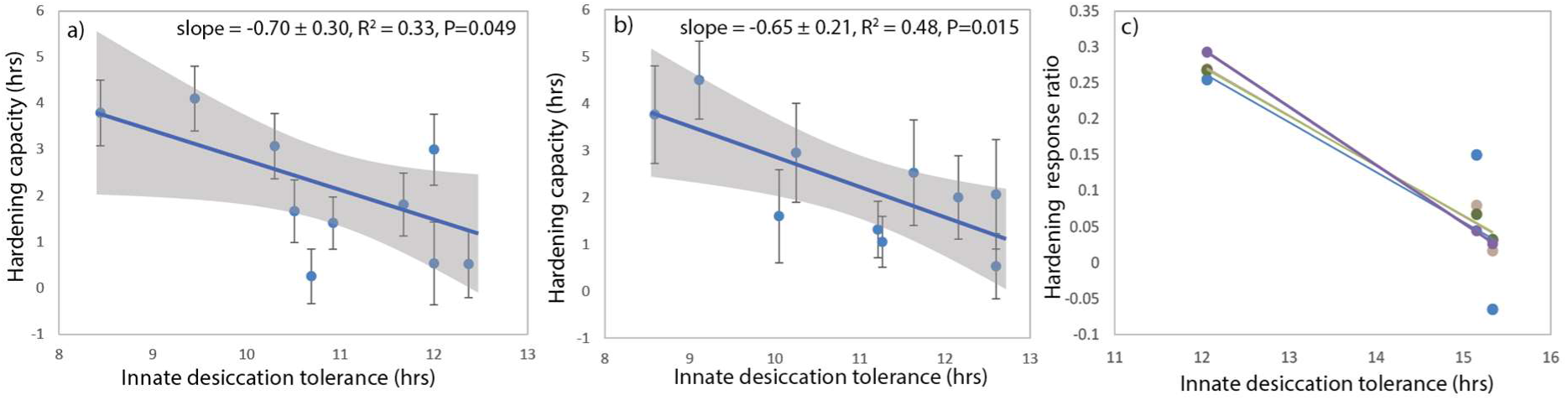
**A)** Relationship between innate desiccation tolerance and hardening capacity. **B)** Relationship between innate desiccation tolerance and hardening capacity, where innate tolerance has been calculated independent from hardening capacity. **C)** Relationship between innate desiccation tolerance and hardening capacity averaged across multiple hardening treatments for three populations

## Discussion

Given that phenotypic plasticity in key traits linked to species distributions is likely to play a vital role in how species respond to climate change, understanding the capacity of different populations to respond via phenotypic plasticity is needed. The two main hypotheses that have been proposed to explain variation in plasticity, the climatic variability and trade-off hypothesis, predict that plasticity is under climatic selection and that phenotypic plasticity does not evolve independently from innate tolerance, respectively. Despite observing variation in plasticity across populations, we found no evidence that climate drives the evolution of hardening capacity in desiccation tolerance. Instead, we show evidence for a trade-off between innate tolerance and plastic capacity across populations.

We found no support for the climatic variability hypothesis in the current dataset. Like other studies, variation in innate desiccation tolerance was linked to climate/latitude (Karan et al. 1998, Parkash et al. 2010, Lasne et al. 2018), suggesting that climate-driven selection is acting on innate tolerance. It should be noted that we did not find a relationship between desiccation tolerance and latitude, unlike Lasne et al. (2008), but this result is in line with other studies along the east-coast of Australia (Hoffmann et al. 2001, Hoffmann et al. 2005). While innate desiccation tolerance was associated with annual precipitation, there was no evidence for climatic selection on phenotypic plasticity. This result supports other studies across a range of traits, finding little evidence for the climatic variability hypothesis across populations (Kelley et al. 2011, Weldon et al. 2018, Healy et al. 2019, Barley et al. 2021). One reason we might see an absence of empirical support for the climatic variability hypothesis is that our estimates of climatic variability do not reflect exposure at the organismal level. Broad-scale climate predictors like WorldClim may not capture the micro-climatic variation that *Drosophila* likely experiences (Pincebourde and Woods 2020). Nevertheless, these broad-scale climate predictors likely correlate with aspects of the environment related to micro-climate variation and the capacity to regulate their exposure to environmental stressors via behaviour. For example, annual precipitation is an important predictor of vegetation cover and, therefore, the capacity to find humid micro-climates (Greve et al. 2011, Hutley et al. 2011). The association between climate and innate tolerance (albeit weak) also suggests that the WorldClim data captures elements of environmental selection. Predictability in environmental variability is also essential for the evolution of phenotypic plasticity via the climatic variability hypothesis (Gabriel and Lynch 1992). The scale of predictability needed for phenotypic plasticity to evolve will depend on an organism’s life history, i.e. how the environment changes within and between generations (Reed et al. 2010). This could partly explain the lack of consistent patterns for the climatic variability hypothesis across species with different life-histories.

Another reason why we might not find evidence for the climatic variability hypothesis is because innate tolerance trade-offs with phenotypic plasticity (van Heerwaarden et al 2024, Stillman 2003; Gunderson and Stillman 2015). This would mean that species that occupy the most extreme climates and are likely to experience the strongest selection for tolerance would have the lowest plasticity, even if they also experienced high climate variability. We found a strong negative correlation when we looked for an association between tolerance and hardening capacity across populations, suggesting a trade-off between these traits. We do not believe this pattern is due to threshold shifts, where tolerant populations require more intense hardening treatments to trigger the highest hardening response (van Heerwaarden et al. 2024), given that our preliminary assessment found no evidence that plasticity thresholds differed across three populations with different levels of innate desiccation tolerance. While we only explored threshold shifts in three populations, given that they varied in their levels of innate desiccation tolerance—ranging from low to high—it is unlikely that threshold shifts are driving trade-off patterns observed here. No differences in plasticity thresholds across populations are similar to patterns overserved in *D. melanogaster* desiccation selection lines, where longer hardening treatments did not increase hardening capacity in the lines selected for higher desiccation tolerance (Hoffmann 1990). However, this result contrasts with evidence for threshold shifts for desiccation and heat tolerance across *Drosophila* species, where species with higher levels of innate tolerance required longer hardening treatments (van Heerwaarden et al. 2024). While it is not clear why threshold shifts are evident across species but not populations, threshold shifts may be more apparent at the species level due to larger differences in innate tolerance in species—ranging from 4 to 37 hours across species, compared to 9 to 13 hours across populations within a species.

As we have previously demonstrated, looking for trade-off patterns across populations (or species) is difficult because of the statistical association between tolerance and plasticity (van Heerwaarden and Kellermann 2020). A common problem in trade-off studies is that innate tolerance is used to assess tolerance and calculate plasticity, and correlating the two defies the assumption of statistical independence. We overcame this issue by creating an alternative data set that calculated innate tolerance independently from hardening capacity (Gunderson 2023). This method reduced our replication, but we still found a significant correlation between innate and hardened capacity. Even so, it is still challenging to know whether trade-offs are underpinned by patterns of selection or physiological/genetic correlations between tolerance and plasticity, with the latter being more important for constraining the evolution of plasticity (van Heerwaarden and Kellermann 2020). Directly selecting on innate and hardened tolerance and then looking at correlated responses in plasticity (while also considering threshold shifts) can establish whether trade-off patterns between tolerance and plasticity are underpinned by pleiotropy or an upper physiological thermal limit. A study comparing hardening responses in *D. melanogaster* lines selected for high desiccation tolerance did find that selected lines showed a 4% increase in desiccation tolerance after hardening, compared to a 24% increase in the unselected lines, in line with an upper physiological limit (Hoffmann 1990).

Although the climatic variability and trade-off hypotheses have been proposed to influence the evolution of plasticity and explain differences in plasticity across organisms, there is mixed evidence for both. By exploring intra-specific variation in tolerance and plasticity across a clinal gradient, we could look at the relative contribution of each to local adaptation in tolerance and plasticity. While innate desiccation tolerance showed evidence of local adaptation, increasing with decreasing precipitation, neither hardened desiccation tolerance or plastic capacity was associated with precipitation or climate variability. Instead, plasticity was associated with basal desiccation tolerance and was lower in populations with high tolerance. These results suggest that innate tolerance was a better predictor of variation in plasticity across populations than climate variability. Together with patterns across species (Kellermann et al. 2018, van Heerwaarden et al. 2024) and from selection studies (Hoffmann 1990), this suggests that tolerance:plasticity trade-offs may often constrain local adaptation in plasticity for desiccation tolerance in *Drosophila*. We encourage future studies to explore trade-offs at different levels.

## Acknowledgements

We acknowledge Shaun Blackburn, Fiona Cockerell, Richard Foo Heng Lee and Allannah Clemson for technical support, and Carla Sgro for comments on an earlier version of the manuscript. Funding. We would like to thank the Australian Research Council for financial support to B.V.H., and V.M.K. through their Fellowship schemes (FT200100025 and FT200100703)

## Notes

### Competing Interest Statement

The authors have declared no competing interest.

## References

Barley, J. M., B. S. Cheng, M. Sasaki, S. Gignoux-Wolfsohn, C. G. Hays, A. B. Putnam, S. Sheth, A. R. Villeneuve, and M. Kelly. 2021. Limited plasticity in thermally tolerant ectotherm populations: evidence for a trade-off. Proceedings of the Royal Society B: Biological Sciences 288:20210765.

Chevin, L. M., and A. A. Hoffmann. 2017. Evolution of phenotypic plasticity in extreme environments. Philosophical Transactions of the Royal Society B-Biological Sciences 372:12.

Chevin, L. M., R. Lande, and G. M. Mace. 2010. Adaptation, plasticity, and extinction in a changing environment: towards a predictive theory. PLoS Biology 8: e1000357.

Chown, S. L., J. G. Sorensen, and J. S. Terblanche. 2011. Water loss in insects: An environmental change perspective. Journal of Insect Physiology 57:1070–1084.

Clemson, A. S., C. M. Sgrò, and M. Telonis-Scott. 2018. Transcriptional profiles of plasticity for desiccation stress in *Drosophil*a. Comparative Biochemistry and Physiology B-Biochemistry & Molecular Biology 216:1–9.

Gabriel, W., and M. Lynch. 1992. The selective advantage of reaction norms for environmental tolerance. Journal of Evolutionary Biology 5:41–59.

Ghalambor, C. K., R. B. Huey, P. R. Martin, J. J. Tewksbury, and G. Wang. 2006. Are mountain passes higher in the tropics? Janzen’s hypothesis revisited. Integrative and Comparative Biology 46:5–17.

Greve, M., A. M. Lykke, A. Blach-Overgaard, and J. C. Svenning. 2011. Environmental and anthropogenic determinants of vegetation distribution across Africa. Global Ecology and Biogeography 20:661–674.

Gunderson, A. R. 2023. Trade-offs between baseline thermal tolerance and thermal tolerance plasticity are much less common than it appears. Global Change Biology 29:3519–3524.

Gunderson, A. R., and J. H. Stillman. 2015. Plasticity in thermal tolerance has limited potential to buffer ectotherms from global warming. Proceedings of the Royal Society B-Biological Sciences 282: 20150401.

Healy, T. M., A. K. Bock, and R. S. Burton. 2019. Variation in developmental temperature alters adulthood plasticity of thermal tolerance in *Tigriopus californicus*. Journal of Experimental Biology 222: jeb213405.

Hoffmann, A. A. 1990. Acclimation for desiccation resistance in *Drosophila melanogaster* and the assocation between acclimation responses and genetic variation. Journal of Insect Physiology 36:885–891.

Hoffmann, A. A., R. Hallas, C. Sinclair, and P. Mitrovski. 2001. Levels of variation in stress resistance in *Drosophila* among strains, local populations, and geographic regions: Patterns for desiccation, starvation, cold resistance, and associated traits. Evolution 55:1621–1630.

Hoffmann, A. A., J. Shirriffs, and M. Scott. 2005. Relative importance of plastic vs genetic factors in adaptive differentiation: geographical variation for stress resistance in *Drosophila melanogaster* from eastern Australia. Functional Ecology 19:222–227.

Hoffmann, A. A., J. G. Sorensen, and V. Loeschcke. 2003. Adaptation of *Drosophila* to temperature extremes: bringing together quantitative and molecular approaches. Journal of Thermal Biology 28:175–216.

Hutley, L. B., J. Beringer, P. R. Isaac, J. M. Hacker, and L. A. Cernusak. 2011. A sub-continental scale living laboratory: Spatial patterns of savanna vegetation over a rainfall gradient in northern Australia. Agricultural and Forest Meteorology 151:1417–1428.

Janzen, D. H. 1967. Why mountain passes are higher in tropics. American Naturalist 101:233–249.

Jensen, A., T. Alemu, T. Alemneh, C. Pertoldi, and S. Bahrndorff. 2019. Thermal acclimation and adaptation across populations in a broadly distributed soil arthropod. Functional Ecology 33:833–845.

Karan, D., N. Dahiya, A. K. Munjal, P. Gibert, B. Moreteau, R. Parkash, and J. R. David. 1998. Desiccation and starvation tolerance of adult *Drosophila*: Opposite latitudinal clines in natural populations of three different species. Evolution 52:825–831.

Kellermann, V., A. A. Hoffmann, J. Overgaard, V. Loeschcke, and C. M. Sgro. 2018. Plasticity for desiccation tolerance across *Drosophila* species is affected by phylogeny and climate in complex ways. Proceedings of the Royal Society B-Biological Sciences 285:20180048.

Kellermann, V., V. Loeschcke, A. A. Hoffmann, T. N. Kristensen, C. Fløjgaard, J. R. David, and J. Overgaard. 2012. Phylogenetic constraints in key functional traits behind species’ climate niches: patterns of desiccation and cold resistance across 95 *Drosophila* species. Evolution 66:3377.

Kellermann, V., and B. van Heerwaarden. 2019. Terrestrial insects and climate change: adaptive responses in key traits. Physiological Entomology 44:99–115.

Kelley, A. L., C. E. de Rivera, and B. A. Buckley. 2011. Intraspecific variation in thermotolerance and morphology of the invasive European green crab, *Carcinus maenas*, on the west coast of North America. Journal of Experimental Marine Biology and Ecology 409:70–78.

Kelly, C., and T. D. Price. 2005. Correcting for regression to the mean in behavior and ecology. American Naturalist 166:700–707.

Lasne, C., S. B. Hangartner, T. Connallon, and C. M. Soro. 2018. Cross-sex genetic correlations and the evolution of sex-specific local adaptation: Insights from classical trait clines in *Drosophila* melanogaster. Evolution 72:1317–1327.

Parkash, R., B. Kalra, and V. Sharma. 2010. Impact of body melanisation on contrasting levels of desiccation resistance in a circumtropical and a generalist Drosophila species. Evolutionary Ecology 24:207–225.

Pereira, R. J., C. M. Saski, and R. S. Burton. 2017. Adaptation to a latitudinal thermal gradient within a widespread copepod species: the contributions of genetic divergence and phenotypic plasticity. Proceedings of the Royal Society B-Biological Sciences 284:20170236.

Pincebourde, S., and H. A. Woods. 2020. There is plenty of room at the bottom: microclimates drive insect vulnerability to climate change. Current Opinion in Insect Science 41:63–70.

Reed, T. E., R. S. Waples, D. E. Schindler, J. J. Hard, and M. T. Kinnison. 2010. Phenotypic plasticity and population viability: the importance of environmental predictability. Proceedings of the Royal Society B-Biological Sciences 277:3391–3400.

Roff, D. A., and D. J. Fairbairn. 2007. The evolution of trade-offs: where are we? Journal of Evolutionary Biology 20:433–447.

Schilthuizen, M., and V. Kellermann. 2014. Contemporary climate change and terrestrial invertebrates: evolutionary versus plastic changes. Evolutionary Applications 7:56–67.

Sgro, C. M., J. Overgaard, T. N. Kristensen, K. A. Mitchell, F. E. Cockerell, and A. A. Hoffmann. 2010. A comprehensive assessment of geographic variation in heat tolerance and hardening capacity in populations of *Drosophila melanogaster* from eastern Australia. Journal of Evolutionary Biology 23:2484–2493.

Sgro, C. M., J. S. Terblanche, and A. A. Hoffmann. 2016. What can plasticity contribute to insect responses to climate change? Annual Review of Entomology 61:433–51

Sinclair, B. J., C. M. Williams, and J. S. Terblanche. 2012. Variation in thermal performance among insect populations. Physiological and Biochemical Zoology 85:594–606.

Sorensen, J. G., T. N. Kristensen, and J. Overgaard. 2016. Evolutionary and ecological patterns of thermal acclimation capacity in *Drosophila*: is it important for keeping up with climate change? Current Opinion in Insect Science 17:98–104.

Sorensen, J. G., and V. Loeschcke. 2001. Larval crowding in *Drosophila melanogaster* induces Hsp70 expression, and leads to increased adult longevity and adult thermal stress resistance. Journal of Insect Physiology 47:1301–1307.

Stillman, J. H. 2003. Acclimation capacity underlies susceptibility to climate change. Science 301:65–65.

Team, R. C. 2014. R: A language and environment for statistical computing.

van Heerwaarden, B., and V. Kellermann. 2020. Does plasticity trade off with basal heat tolerance? Trends in Ecology & Evolution 35:874–885.

van Heerwaarden, B., V. Kellermann, and C. M. Sgro. 2016. Limited scope for plasticity to increase upper thermal limits. Functional Ecology:1947–1956.

van Heerwaarden, B., R. F. H. Lee, J. Overgaard, and C. M. Sgro. 2014. No patterns in thermal plasticity along a latitudinal gradient in *Drosophila simulans* from eastern Australia. Journal of Evolutionary Biology 27:2541–2553.

van Heerwaarden, B., C. Sgrò, and V. M. Kellermann. 2024. Threshold shifts and developmental temperature impact trade-offs between tolerance and plasticity. Proceedings of the Royal Society B: Biological Sciences 291:20232700.

van Noordwijk, A. J., and G. de Jong. 1986. Acquisition and allocation of resources: their influence on variation in life history tactics. The American Naturalist 128:137–142.

Weldon, C. W., C. Nyamukondiwa, M. Karsten, S. L. Chown, and J. S. Terblanche. 2018. Geographic variation and plasticity in climate stress resistance among southern African populations of *Ceratitis capitata* (Wiedemann) (Diptera: Tephritidae). Scientific Reports 8:9849.

