## Supplementary material for "Variation in phenotypic plasticity in desiccation tolerance is driven by trade-offs, not climate, in *Drosophila melanogaster*": Supp

Table S1. Longitude and latitude for the ten sites where we collected Drosophila melanogaster and the nearest town to that collection site

| Site | GPS co-ordinates |
|---|---|
| Townsville | 146.79, -19.26 |
| Rockhampton | 150.72, -23.15 |
| Brisbane | 153.30, -27.61 |
| Coffs Harbour | 153.15, -30.23 |
| Port Macquarie | 152.90, -30.93 |
| Gosford | 151.20, -33.31 |
| Woolongong | 150.91, -34.34 |
| Bermugi | 150.06, -36.40 |
| Melbourne | 145.27, -37.99 |

Table S2. Mixed effects analysis of variance showing the relationship between no exposure to pre-treatment of desiccation stress vs pre-treatment (Treatment) and latitude.

| | D.F. | Chisq | R2m | R2c |
|---|---|---|---|---|
| Treatment | 1 | 3.11 | 0.16 | 0.22 |
| Latitude | 1 | 2.35 | | |
| Tank | 2 | 40.73*** | | |
| Latitude x Treatment | 1 | 0.02 | | |

Table S3. Mixed effects analysis of variance showing the relationship between no exposure to pre-treatment of desiccation stress vs pre-treatment (Treatment) and precipitation seasonality.

| | D.F. | Chisq | R2m | R2c |
|---|---|---|---|---|
| Treatment | 1 | 30.02*** | 0.16 | 0.22 |
| Precipitation seasonality | 1 | 2.35 | | |
| Tank | 2 | 40.82*** | | |
| Precipitation seasonality x Treatment | 1 | 0.24 | | |

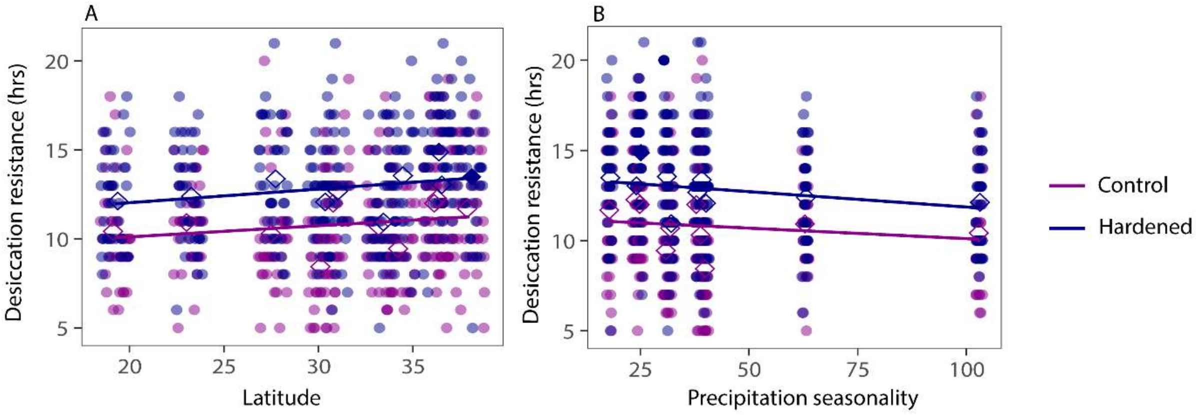


Figure S1. A) Showing the relationship between desiccation tolerance and latitude for hardened and innate desiccation tolerance. B) Showing the relationship between desiccation tolerance and precipitation seasonality for hardened and innate desiccation tolerance. Open diamonds show the population means.
